# Using machine learning to count Antarctic shag (*Leucocarbo bransfieldensis*) nests on images captured by Remotely Piloted Aircraft Systems

**DOI:** 10.1101/2024.02.27.582379

**Authors:** Andrew Cusick, Katarzyna Fudala, Piotr Pasza Storożenko, Jędrzej Świeżewski, Joanna Kaleta, W. Chris Oosthuizen, Christian Pfeifer, Robert Józef Bialik

## Abstract

Using 51 orthomosaics of 11 breeding locations of the Antarctic shag, we propose a method for automating counting of shag nests. This is achieved by training an object detection model based on the YOLO architecture and identifying nests on sections of the orthomosaic, which are later combined with predictions for the entire orthomosaic. Our results show that the current use of Remotely Piloted Aircraft Systems (RPAS) to collect images of areas with shag colonies, combined with machine learning algorithms, can provide reliable and fast estimates of shag nest counts (F1 score > 0.95). By using data from only two shag colonies for training, we show that models can be obtained that generalise well to images of both spatially and temporally distinct colonies. The proposed practical application opens the possibility of using aerial imagery to perform large-scale surveys of Antarctic islands in search of undiscovered shag colonies. We discuss the conditions for optimal performance of the model as well as its limitations. The code, data and trained model allowing for full reproducibility of the results are available at https://github.com/Appsilon/Antarctic-nests.

## INTRODUCTION

In recent years, Remotely Piloted Aircraft Systems (RPAS) have become common tools for wildlife monitoring (Schad and Fisher, 2023) and have proved especially helpful for surveying habitats that are otherwise difficult to access, such as many areas of Antarctica (Pina and Vieira, 2022). RPAS are used to monitor many different wildlife species in the Antarctic environment, including whales, surface-nesting birds and pinnipeds (Fudala and Bialik 2022; Tovar-Sánchez et al. 2021; Zmarz et al. 2018). Long-term monitoring of wildlife populations with RPAS often generate extensive datasets, frequently high-resolution images that are used to build orthomosaics on which animals (or other features) can be counted. Manual counting of individuals on an orthomosaic requires a large investment of researchers’ time. To minimize counting errors and to obtain a measure of repeatability, counting is often carried out by more than one person, thus multiplying the effort needed. Machine learning algorithms provide a way to automate detection and counting of objects in images, increasing the efficiency of wildlife monitoring with RPAS. Computer vision models have, for example, been used to detect seabird guano in satellite images (Witharana and Lynch 2016; Le et al. 2022) and to detect penguins, albatrosses and many other seabird species in images obtained from RPAS (Liu et al. 2020). However, while the use of RPAS in ecological studies is expanding rapidly, the use of machine learning to survey visual data is increasing at a much slower (~50%) rate (Dujon and Schofield 2019).

Antarctic shags (*Leucocarbo bransfieldensis*) is a surface-nesting species that breed in the Antarctic Peninsula and surrounding island groups on islands around Antarctica. Antarctic shags often breed in locations that are completely inaccessible to ground-based surveys (e.g., sea cliffs, islets or elevated capes), making aerial photography the only reliable means available to survey some breeding sites. *L. bransfieldensis* were a trigger species for the establishment of 24 of 205 Antarctic Important Bird and Biodiversity Areas (IBAs). Monitoring their population trends are therefore important (Schrimpf et al. 2018), but to our knowledge, there is currently no method available to automatically detect and count Antarctic shags in RPAS images.

In this study, we use an object detection machine learning model to identify and count individual Antarctic shag nests in georeferenced aerial images collected by RPAS. We chose to count individual nests (a proxy of breeding pairs) rather than individual birds because Antarctic shags and chinstrap penguins sometimes breed in mixed colonies. Antarctic shags have a white patch of plumage on their backs, seen on aerial imagery as a “white spot on a black background", which can be used to distinguish shags from penguins. However, the white spot on the back is only visible in when birds sit in certain postures, and the similar body dimensions of shags and chinstrap penguins, therefore necessitate the use of more than one distinguishing feature where these species occupy a common space. Antarctic shag nests are three-dimensional objects that are built of organic material, mainly guano-cemented marine algae and feathers (Bernstein and Maxson, 1982). These nests are highly recognizable on aerial photographs even though their appearance may take different forms through the breeding season. The diameter of shag nests is approximately 49 cm (SD = 5 cm, n = 43) (Pfeifer et al., 2021), which is more than 50 times larger than the image pixel that can be obtained using RPAS when operating at recommended RPAS flying altitudes in Antarctica (Weimerskirch et al. 2017; Harris et al. 2019). Hence, nests are ideal objects for counting on aerial images. We collected 51 RPAS orthomosaics from 11 breeding locations of Antarctic shags and tested whether an object detection machine learning model trained on images from two colonies could accurately count nests separated in time (at the training colonies), or in space (at other colonies). We discuss the performance of the model to count nests and identify colony locations, and the benefits and limitations of applying this method.

## DATA AND METHODS

### Orthomosaics

The data used in the study were collected by three independent research groups using various types of RPAS (Table 1). In total, 51 surveys of 11 sites around King George Island and Nelson Island on the South Shetland Islands were flown (Figure 1). For all the locations, there were multiple images and we used Pix4Dmapper (Pix4D S.A., Prilly, Switzerland) to generate a single orthomosaic of the area of interest. Notably, the orthomosaics varied in spatial resolution and visual characteristics (level of focus, localised spatial distortions, chromatic aberrations, etc.). Details on the size and characteristics of the data used in the study are presented in Table S1.

**Table 1.**
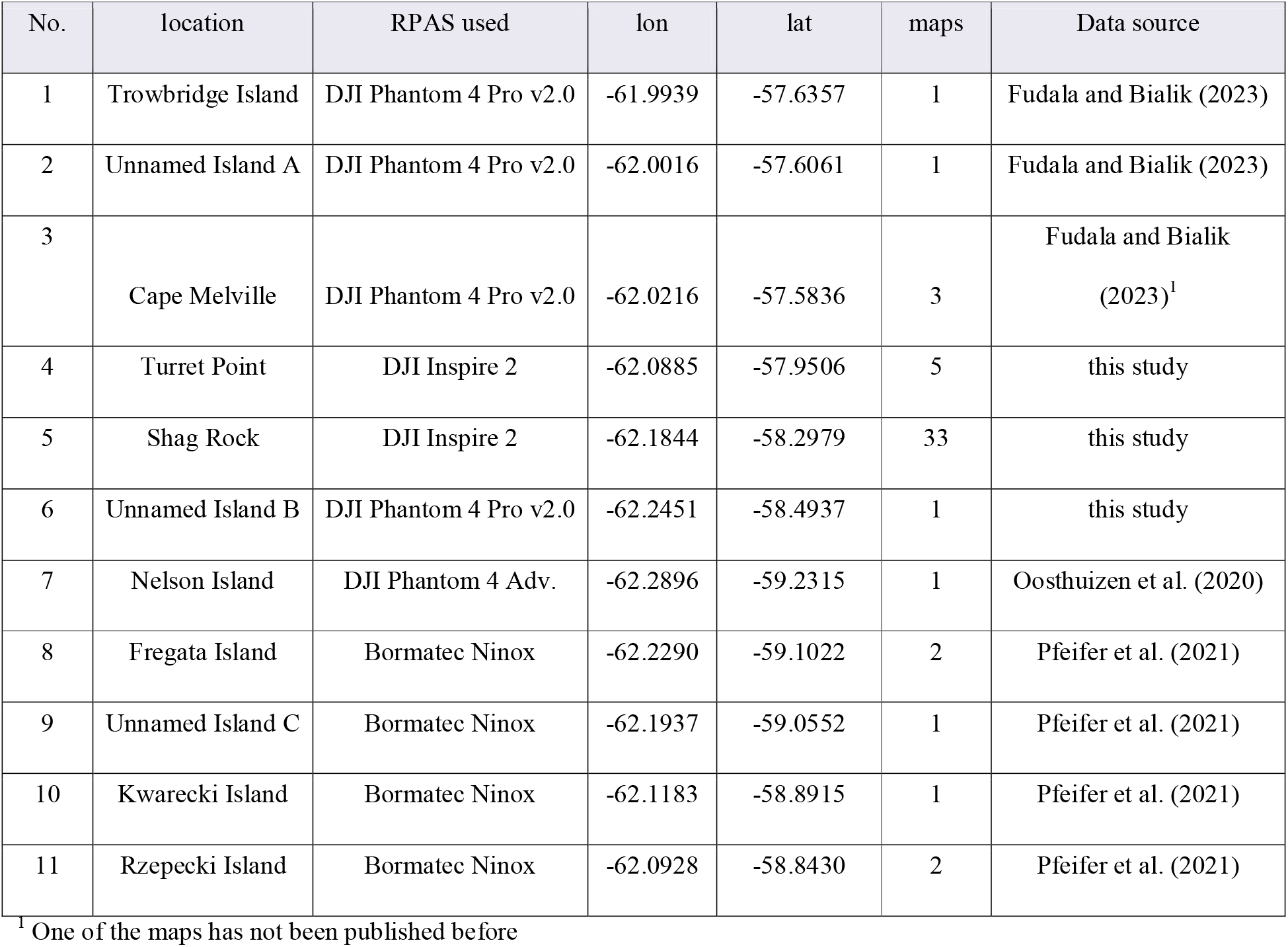
Details of the photogrammetry missions. The sequence numbers correspond to the locations in Figure 1.

| No. | location | RPAS used | lon | lat | maps | Data source |
| --- | --- | --- | --- | --- | --- | --- |
| 1 | Trowbridge Island | DJI Phantom 4 Pro v2.0 | -61.9939 | -57.6357 | 1 | Fudala and Bialik (2023) |
| 2 | Unnamed Island A | DJI Phantom 4 Pro v2.0 | -62.0016 | -57.6061 | 1 | Fudala and Bialik (2023) |
| 3 | Cape Melville | DJI Phantom 4 Pro v2.0 | -62.0216 | -57.5836 | 3 | Fudala and Bialik (2023)<sup>1</sup> |
| 4 | Turret Point | DJI Inspire 2 | -62.0885 | -57.9506 | 5 | this study |
| 5 | Shag Rock | DJI Inspire 2 | -62.1844 | -58.2979 | 33 | this study |
| 6 | Unnamed Island B | DJI Phantom 4 Pro v2.0 | -62.2451 | -58.4937 | 1 | this study |
| 7 | Nelson Island | DJI Phantom 4 Adv. | -62.2896 | -59.2315 | 1 | Oosthuizen et al. (2020) |
| 8 | Fregata Island | Bormatec Ninox | -62.2290 | -59.1022 | 2 | Pfeifer et al. (2021) |
| 9 | Unnamed Island C | Bormatec Ninox | -62.1937 | -59.0552 | 1 | Pfeifer et al. (2021) |
| 10 | Kwarecki Island | Bormatec Ninox | -62.1183 | -58.8915 | 1 | Pfeifer et al. (2021) |
| 11 | Rzepecki Island | Bormatec Ninox | -62.0928 | -58.8430 | 2 | Pfeifer et al. (2021) |

<sup>1</sup> One of the maps has not been published before

**Fig. 1.**
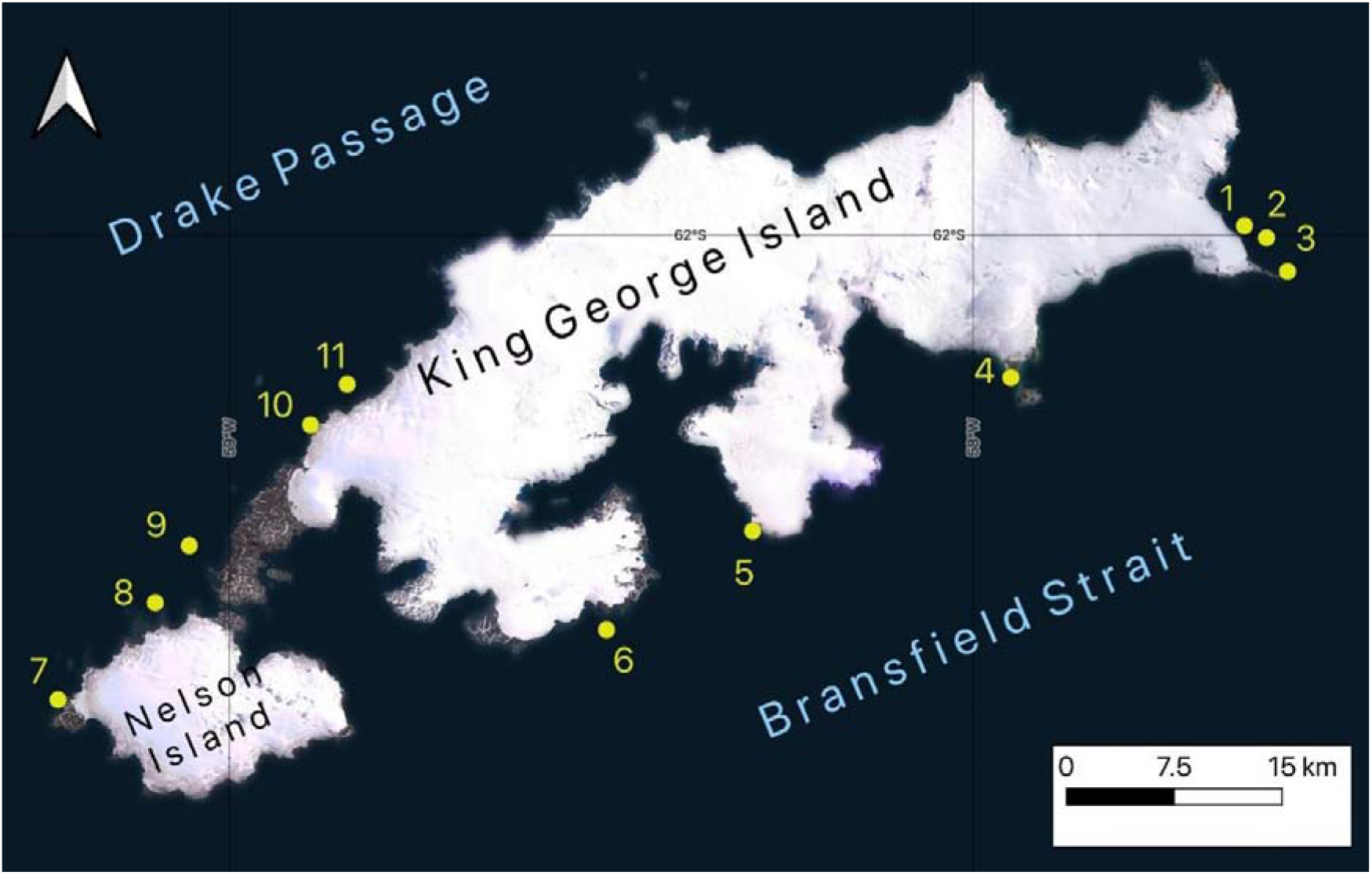
Locations of Antarctic shag colonies considered in the study. The study colonies (1 to 11) are listed in Table 1.

### Labels

We visually identified all the shag nests in each orthomosaic and assigned labels to the centre of each nest via QGIS software (QGIS 3.16.5 ‘Hannover’). When shags and penguins were breeding in close proximity (Figure 2), we used additional supporting imagery taken from the ground or expert knowledge of the particular location to assist labelling.

**Fig. 2.**
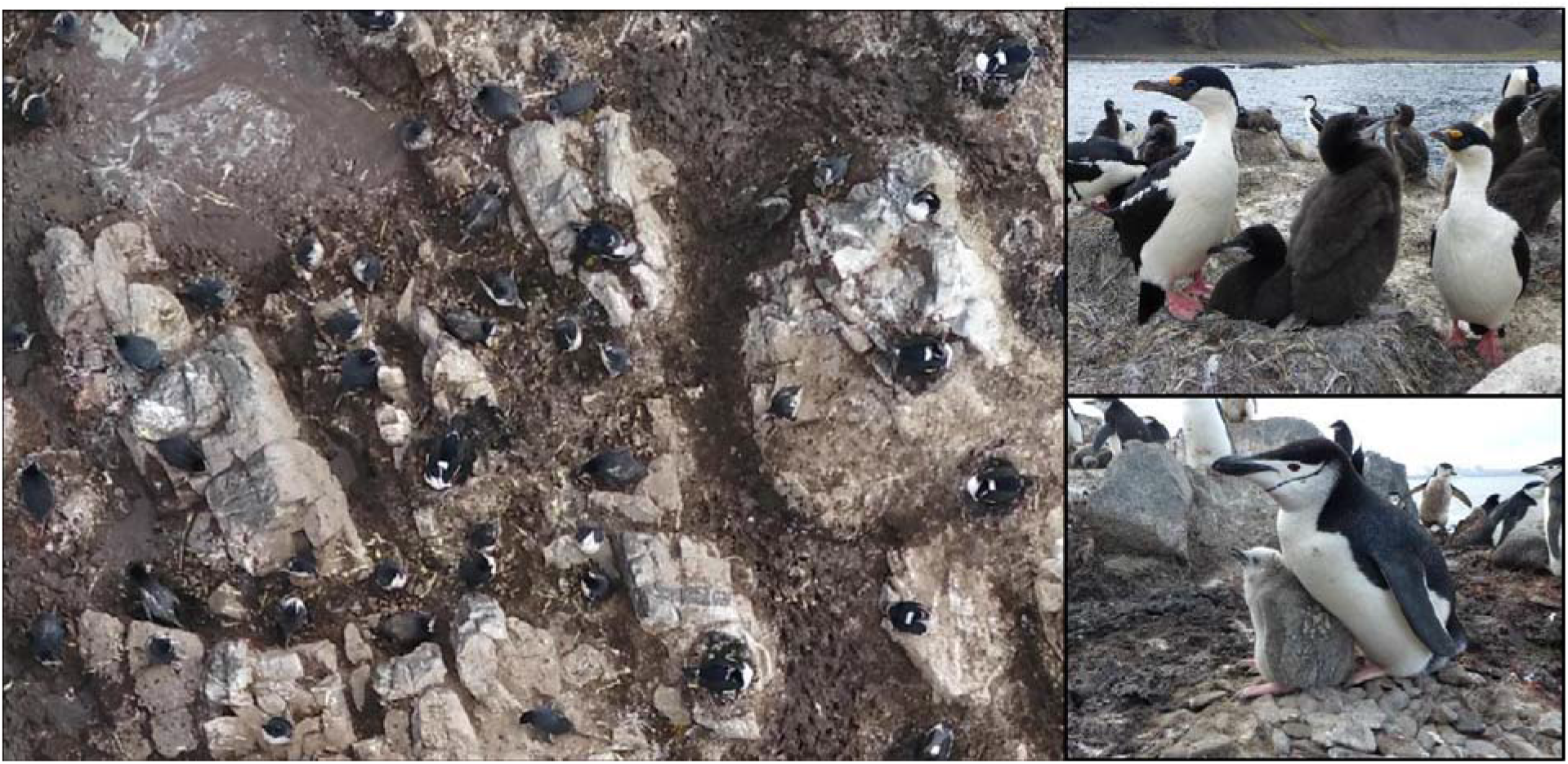
Antarctic shags often breed in mixed colonies with chinstrap penguins. Similar body dimensions and plumage colours can complicate the identification of the two species in aerial photographs.

Antarctic shag nests have a regular, circular shape and are approximately 50 cm in diameter (Pfeifer et al. 2021). Therefore, the locations of the centres of the nests were translated into square bounding boxes of 50 cm × 50 cm. The bounding boxes were defined geographically, but since the images were georeferenced, they could be translated to specific pixels of relevant images (Figure 3).

**Fig. 3.**
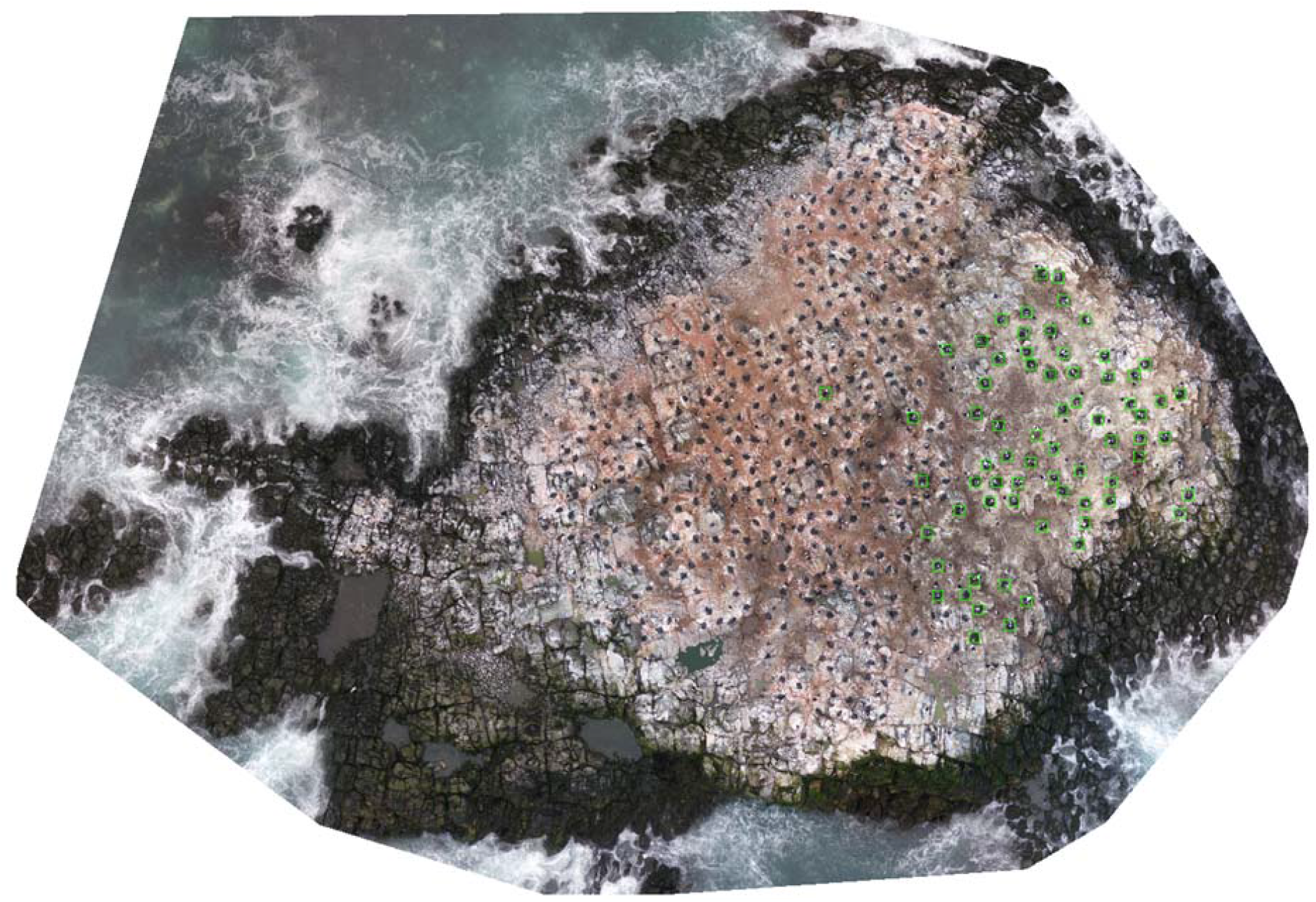
Orthomosaic of the Shag Rock colony with shag nests labelled using 50 cm × 50 cm bounding boxes (in green). Note the presence of some shag’s nest within the penguin colony.

### Dataset creation

The orthomosaics available for the study varied significantly in spatial resolution (from 6 mm to 34 mm per pixel; see Table S1). The images were rescaled to a common resolution of 20 mm per pixel, for two reasons: first, to achieve a model able to perform well also at lower resolutions; second, to leverage the constant size of the nests in terms of occupied pixels. The choice of the resolution used was dictated by the data available for the study (only two orthomosaics had lower resolutions) and the intention to leave at least 25 × 25 pixels per nest. The images used in the study captured significant geographical distances (400 m in the case of Cape Melville), while the objects of interest, i.e., the nests, were only 50 cm in size. Hence, an approach involving tiling (also often referred to as patching) was used. All the orthomosaics were cut into square tiles of 12.8 m × 12.8 m along longitudinal and latitudinal directions, using a custom Python script written for that purpose and available in the shared code repository. In this way, each tile was 640 × 640 pixels. For the orthomosaics that had lower resolution than the one used in the analysis (e.g., Nelson Island and Kwarecki Island), the resulting tiles contained less information than could have been fitted in the tiles, with their pixels stretched to preserve the geographical distances needed. Tiling was performed with 6.4 m overlaps in the longitudinal and latitudinal directions so that the majority of the nests were captured on four individual tiles. The key reason for introducing the overlap and making it so significant is that this way each nest would be fully present in at least one of the tiles.

This approach has two side effects to be considered. First, most of the nests appear multiple times in the data, which requires attention when combining model predictions per image, as they need to be automatically combined into a single prediction per actual nest. To overcome this, non-max suppression is applied after collecting the predictions for a given image (and not for individual tiles). Second, using the overlap increases the number of tiles that need to be analysed by a model at inference time. This is a reasonable trade-off for most practical use cases since real-time results are not needed and therefore speed of analysis is considerably less important than model performance (having more undetected nests).

### Training data

Two locations were chosen to constitute the training dataset: Shag Rock and Cape Melville. For Shag Rock, two images were left out of the training dataset and were instead included in the test set. Those images were captured several months after the last of image from the training dataset from Shag Rock. Similarly, for Cape Melville, two images captured in December 2022 were used for training, while an image taken in November 2023 was used for testing. In this way, the robustness of the model against temporal separation can be tested. To balance the proportion of tiles including instances of nests, we first selected only tiles with nests; from the remaining tiles, we randomly selected a corresponding number of tiles without nests. With this selection, the training dataset spans 2.5 years and includes images of shag nests in various stages of its annual reproductive cycle, including empty nests. With an extensive dataset covering a cross-section of the entire year, it was possible to train the model on a variety of substrate and shape variants on which nests occurred during the period (Figure 4).

**Fig. 4.**
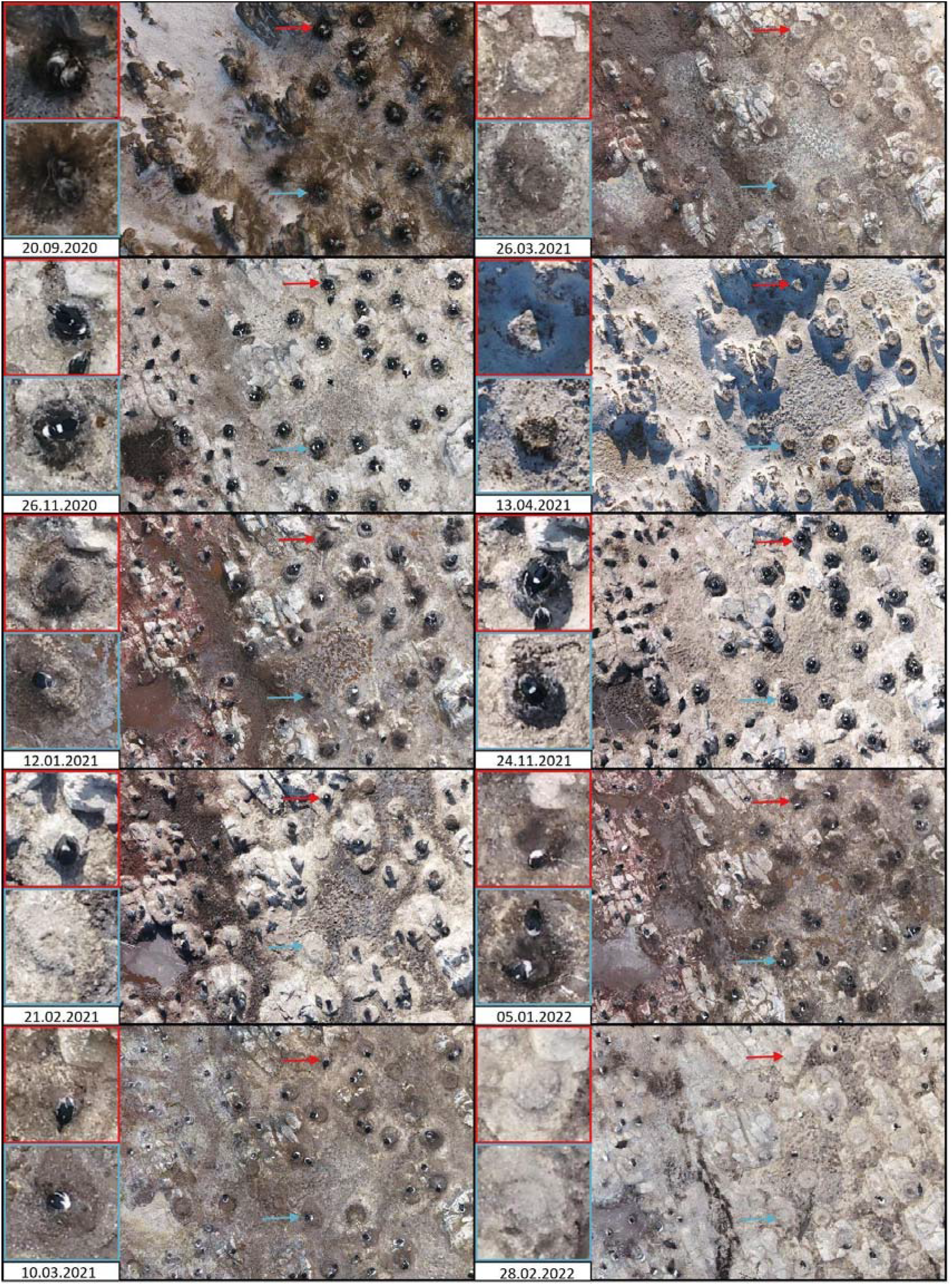
Antarctic shag nests and variation in the colony substrate at different times during the phenological cycle. Red and blue arrows indicate the same nest on 10 different dates.

### Validation data

For validation of the model performance, during the training process (after each training epoch), we used three orthomosaics of Turret Point (with ~2/3 covered by a shag colony), which were spatially distinct from the orthomosaics included in the training dataset (Shag Rock and Cape Melville). Two (temporally distinct) out of five images of Turret Point were left out of the validation set and added to the test set.

### Test data

The performance of the models was tested on the following groups of data:

1. Orthomosaics of Shag Rock, Cape Melville, and Turret Point, temporally distinct from those in the training or validation data (i.e., coming from a consecutive nesting season). Referred to later as “test-over_time”.
2. Orthomosaics of locations geographically distinct from both the training and validation datasets in Unnamed Island A, Unnamed Island B and Trowbridge Island. Referred to later as “test-over_location”.
3. Orthomosaics of locations geographically distinct from both the training and validation datasets (Kwarecki, Rzepecki, Fregata, Nelson and Unnamed Island C) and collected with older types of RPAS and cameras (with imagery and labels provided by external experts). Referred to later as “test-over_source”.

For the test data, the entire set of available images was used for tiling, which in some cases meant significant portions devoid of any nests (e.g., showing water).

## MACHINE LEARNING MODEL

### Model definition

To detect shag nests automatically, several of the YOLO state-of-the-art object detection framework models (YOLOv5, YOLOv6L and YOLOv8) were tested. The model that performed the best was YOLOv6L. We employed the standard architecture of the large YOLOv6 model with the recommended hyperparameters for fine-tuning [https://github.com/meituan/YOLOv6] and standard data augmentation parameters that are largely consistent across YOLO versions. The training time was approximately 10 hours on a single Nvidia T4 GPU. We experimented with using different levels of overlap in both the training data and at inference time. We used many different sets of hyperparameters in YOLOv6 but found only marginal improvements; hence, we decided to use the default settings. Finally, we tested working with different spatial resolutions of the imagery, most notably including training runs in which the same images were used at different spatial resolutions. We did not find this approach to be beneficial.

### Inference method

Object detection models typically return bounding box proposals, which are equipped with scores indicating the confidence level of a given prediction. The following method was used to combine predictions obtained for individual tiles into a collection of final predictions per orthomosaic:

1. Since we cut the data into overlapping tiles and some nests may have landed at the border of the tile, we removed all predictions closer than 50 cm to the border. Nevertheless, all nests at the tile borders are visible on other tiles.
2. To have only one prediction per nest, we removed lower confidence predictions that significantly overlapped with one another (those that had intersection-over-union above the threshold of 0.2, which for equal areas of two predictions means that one-third is common).
3. We filtered the predicted bounding boxes according to their size (requiring that neither side be smaller than 0.3 m or larger than 0.7 m, the area of the nest be larger than 0.125 m^2^ and the ratio of the sides be larger than 0.8).
4. Finally, to assess the counts of the predicted nests, a confidence score threshold of 0.5 was used (notably, this threshold was not used in the calculation of mean average precision (mAP) scores; see Supplement).

### Model evaluation

After the model predictions for individual tiles were combined into a set of predictions for the entire orthomosaic, the quality of the predictions could be assessed. For the purpose of testing the models, we used the tiling of the entire available test orthomosaics. The quantitative assessment was based on an approach tailored to the use case. First, a threshold of confidence was applied to determine the location the model predicts for the nests (e.g., allowing counting of the nests predicted by the model

-a key ecological indicator). Each prediction was considered as a true positive (TP) if there was a nest with which it overlapped significantly (had an intersection over union above the threshold of 0.3). Each nest could be predicted only once. The predictions that were not matched to a nest were treated as false positives (FP), while the nests that were not assigned to any prediction were treated as missed or false negatives (FN). To provide an overview of the results, the F1 score was used. Based on the above definitions, F1 is calculated by the following formula (Van Rijsbergen, 1979):

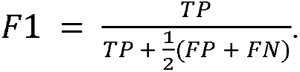

This metric is the harmonic mean between precision and recall and is a commonly used (e.g., Peng et al. 2020; Roy et al. 2023), conservative measure of performance that balances both missing predictions (FN) and not predicting too many instances (FP). When F1 takes its highest value of 1.0, it indicates excellent precision and recall; when precision or recall are zero, the value of F1 is 0.

## RESULTS

### Model performance

The model achieved nearly perfect F1 scores on all the training and validation data (Figure 5). For the testing data, it generalised exceptionally well over time at all locations (“test-over_time”). Moreover, a nearly perfect F1 score was also obtained at locations it had never seen before (“test-over_location”) (Figure 6). In Table S2, we report the F1 scores as well as the number of actual nests, TP, FP and FN for each individual orthomosaic. Notably, the three orthomosaics in test-over_location contained 48 nests. The model correctly detected all of them and only produced one incorrect prediction — predicting that an apparent shag is sitting on a nest. Detailed graphs with each orthomosaic’s score presented separately can be found in the Supplement (Figure S2).

**Fig. 5.**
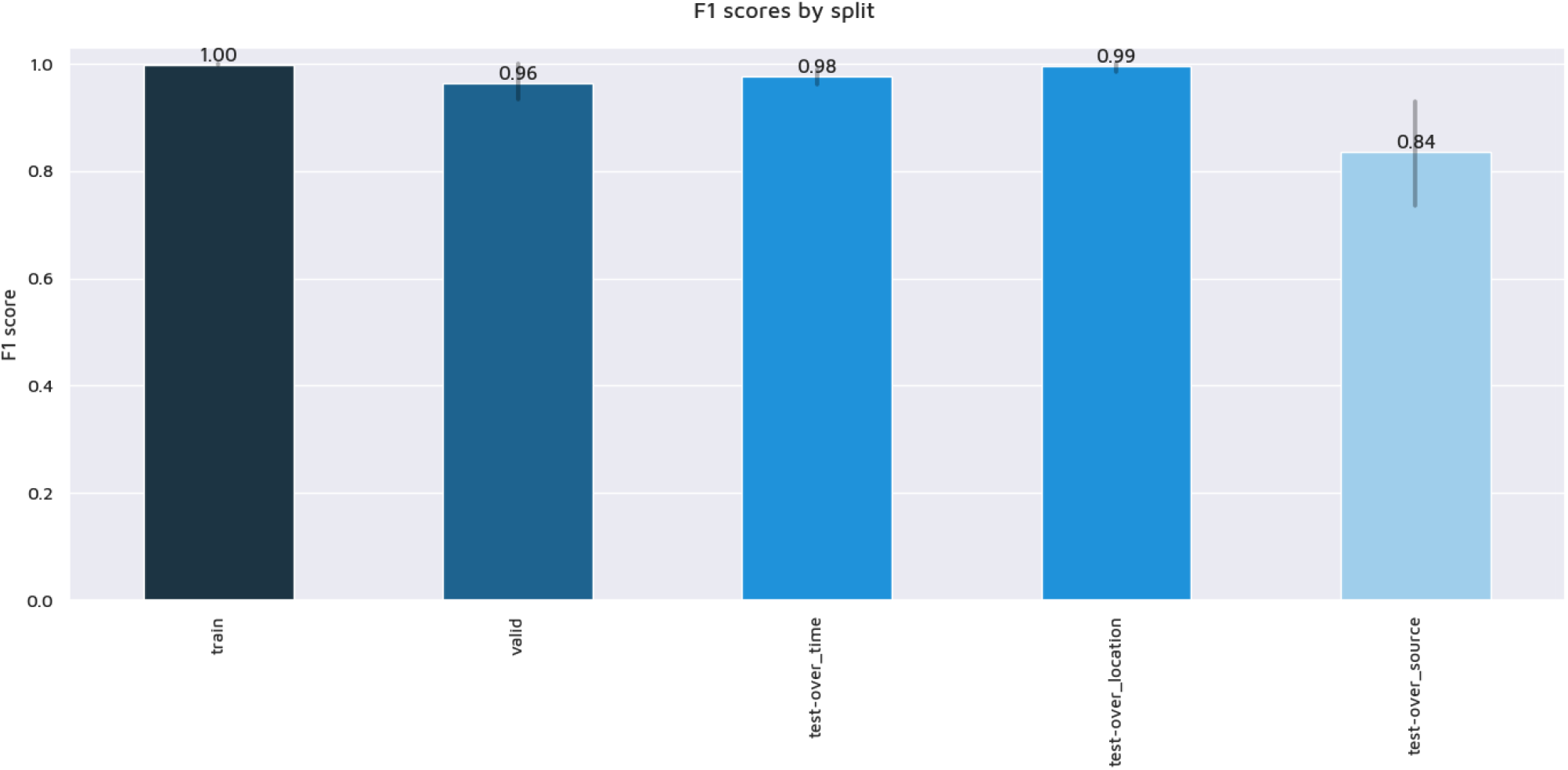
Summary of the F1 scores obtained on different parts of the data. The near perfect score obtained by the model on the training and validation data was also reproduced on the test data. Details regarding the scores for individual orthomosaics can be found in Table S2 and Figure S2.

**Fig. 6.**
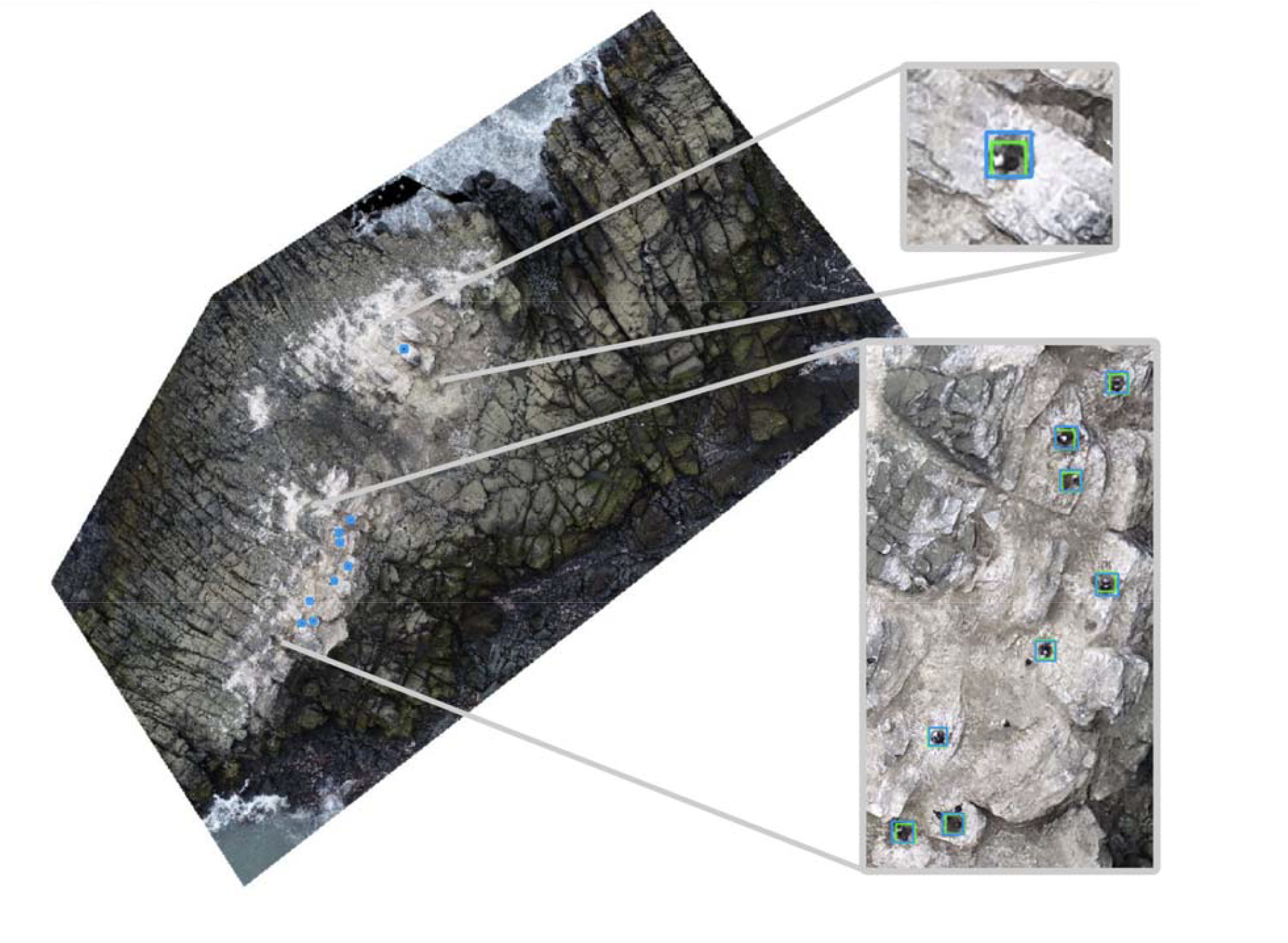
Ground truth nests (the nine green squares) and the predictions made by the model (blue boxes) for a test-over_location orthomosaic of Unnamed Island B. Note that all the nests are correctly located (even the one that is spatially disjointed from the rest), with no false positive predictions.

Even though the orthomosaics often covered large areas, with colonies of shags occupying a small portion of the field of view, false positive predictions of nests almost never happened, while nests distant from the colony were correctly identified. But, it is informative to inspect the few cases where the model made mistakes. The two lowest F1 scores were obtained for Fregata and Rzepecki Islands. In both cases, the orthomosaics had optical artefacts such as out-of-focus blur and colour aberration present in the entire image, and the area with the nests was distorted, blurred and stretched possibly as a result of being located on a hillside (Figure 7). Fortunately, in both cases, we had a second orthomosaic each captured within days from the first one. In both cases, the results of the model on the orthomosaic of better visual quality yielded scores as high as those achieved on the remaining studied data. Notably, the orthomosaic of Rzepecki Island captured on 2016-12-30 received a nearly perfect score, with 58 nests out of 62 identified correctly and no false predictions.

**Fig. 7.**
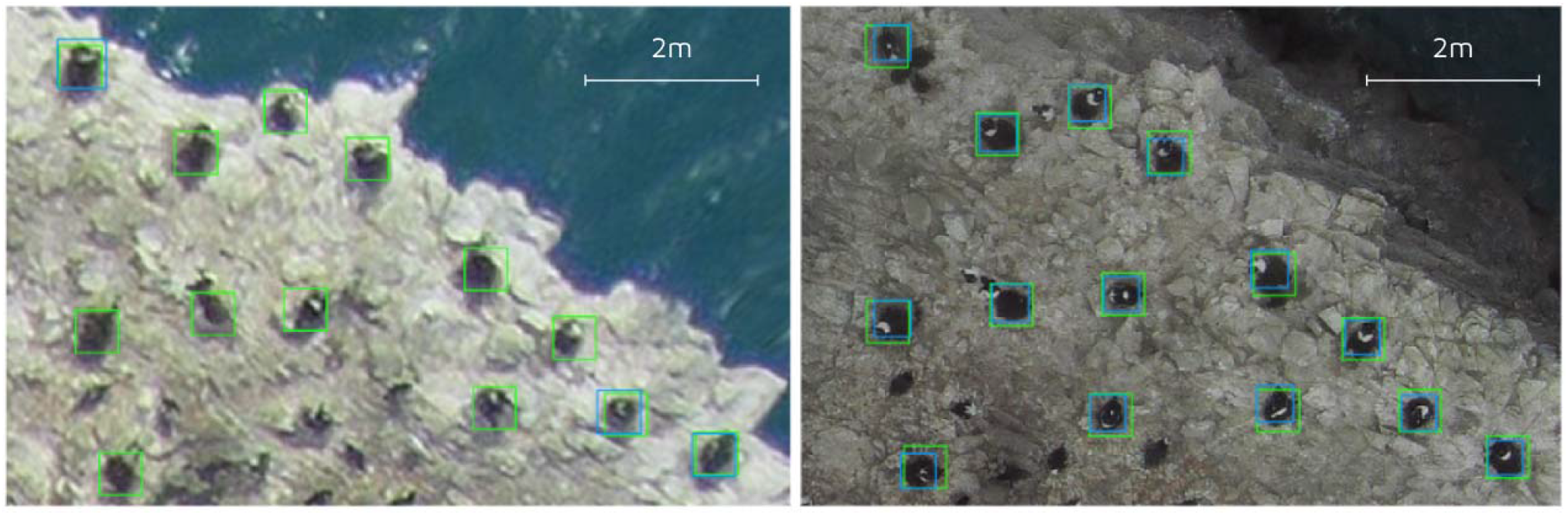
Close-up image of an area of Rzepecki Island from the test-over_source (ground truth nests are marked in green, and predictions are marked in blue). The image in the left panel is blurred, and only three nests from this part of the island were identified by the model. The image in the right panel shows the same area, which was captured two days later. The improved sharpness of the image allows the model to pick up all the nests correctly (even though the resolution is low and the image is significantly darker than the well-lit training images).

Cape Melville is also worth mentioning. While two orthomosaics of this large colony (eight times more nests than the second largest, Shag Rock and Nelson Island) were present in the training data, they were both collected during good weather conditions, with no snow in the colony and the nests clearly visible. The orthomosaic used in the test dataset (collected almost a year later), included large areas covered by snow, which obscured some of the nests. While the F1 score was not ideal for this map (0.94), the number of predicted nests was within a marginal error of 0.8% of the actual number of 478.

### Issues identified in the data

There were several shortcomings in the quality of the imagery used in the study. First, it is common practice to collect images as individual pictures with a fixed set of camera settings and RPAS altitudes. This, in some cases, resulted in images varying in the appearance of the colours and in parts of the images (e.g., containing parts of the islands that were higher than their neighbourhood) being out of focus. Second, the individual images are combined into a single map with noticeable marks on the patching, including in particular the borders of pixels occupied by shags. Since the birds are often occupying their nests, this had a bearing on the visual quality of the objects studied in this study. Third, the data collected in 2016, which were used as part of the test set, contained in some cases significant optical artefacts, such as motion blur, chromatic aberration and spatial distortions (Figure 8).

**Fig. 8.**
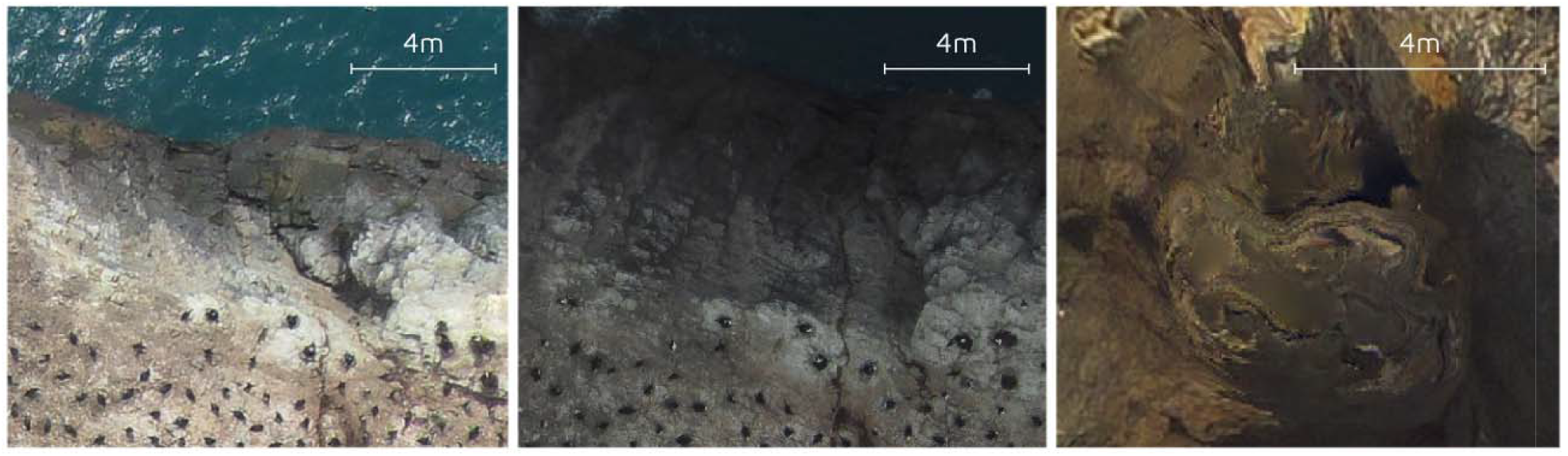
Examples of shortcomings in the quality of the data used in the study. Left and centre show the same area of Rzepecki Island in orthomosaics taken at 2-day intervals; right shows a close-up of distortion on Kwarecki Island.

### Computational requirements

Inference on a single tile took approximately 60 ms using an Nvidia T4 GPU. Inference on tiles from all the orthomosaics in this study took 15 min on the GPU, and they covered an area of 0.6 km^2^. This means that inference for tiles covering an area of 1 km × 1 km, would take approximately 25 min. While the post-processing time of the predictions (merging predictions per tile into a single prediction per image) is negligible, additional time may be required to prepare the tiles. Rescaling and cutting a large orthomosaic into tiles can take significantly longer depending on the starting resolution and hence the size of the image representing the orthomosaic.

## DISCUSSION

The possibility of using RPAS for large-area mapping of locations where Antarctic shags nest seems more feasible now than ever before, particularly using fixed-wing drones (Pfeifer et al. 2021, Zmarz et al. 2018). RPAS surveys coupled with computer vision models can therefore be a valuable tool for ecosystem monitoring. Such approaches offer new possibilities, but automated methods are required for analysing RPAS images or orthomosaics. Our results show that modern machine learning models can provide reliable counts of Antarctic shag nests. Coupling RPAS surveys with machine learning processing of images can thus be used to monitor species that are difficult to count in other ways. Machine learning provides a solution for simplifying the processing of aerial imagery and allowing researchers to more easily, efficiently and accurately extract ecological data from large amounts of imagery data. To make these methods more accessible, we published the data used in this study, the code used in the analysis and the trained machine learning model.

We used F1 scores to evaluate model performance, and in all cases found that the models were able to predict counts with high accuracy. The performance of object detection models is often measured by a standard metric of mean average precision (often referred to as mAP or mAP[0.5:0.95]) (see Eikelboom, et al. 2019, Moreni et al. 2023). We present such an evaluation of the model in the supplement (see Figure S1), but due to its focus on the exact overlap between the ground truth and predicted bounding boxes, we do not believe this metric is informative for the current study. For example, one of the lowest mAP values (0.36) was achieved on Unnamed Island B, for which all the nests were identified and no false positives were produced by the model (see Figure 6). This is because while the model learned to exactly highlight the area of the nests, the ground truth used for the assessment of the model was derived from fixed-size boxes centred on an approximate physical centre of the nests.

In addition to known colonies, our study also shows that RPAS are useful tools to detect new colonies of seabirds. The global population size of the Antarctic shag is estimated to be approximately 12,191 breeding pairs nesting in c. 175 breeding colonies (Fudala and Bialik, 2023) that are often located on rocky slopes (Harrison et al., 2021, Oosthuizen et al., 2020). The need for an evaluation of the status of the population was highlighted by Schrimpf et al. (2018); since then, two new colonies have been inventoried at Ryder Bay (Phillips et al. 2019) and at Cape Melville (Fudala and Bialik, 2023). Orthomosaics of eleven locations where Antarctic shags breed were used in this work. Three of these islands/islets are unnamed (designated Unnamed Islands A-C). The breeding group on Unnamed Island B was identified for the first time in this work. There are hundreds of such islets along the northern parts of Nelson and King George Islands (Pfeifer et al. 2021). Some of them were inventoried decades ago, and the presence of bird colonies needs to be verified, as shown by the example of the location of Smilets Point, where the Antarctic shag nested in 1986/1987 (Shuford and Spear 1988) and no nests were found at this location in 2016/2017 (Pfeifer et al. 2021). Coastal islets inaccessible to terrestrial observers may account for a significant and still severely underestimated share of the total breeding population locations. In the case of the entire range of the species, there could be thousands of such islands/islets.

Most of the orthomosaics used in the study contained large (sometimes very large - reaching tens of thousands of individuals) colonies of penguins. Hence, these orthomosaics can be used for analogous studies of penguin populations. The challenge lies in obtaining accurate annotations for those much larger numbers of objects to identify. We plan to address this in further research.

## ACKNOWLEDGEMENTS

KF and RB were supported by Institute of Biochemistry and Biophysics PAS Internal grant MG-09/2021 and by the Ministry of Science and Higher Education of Poland grant no. 6812/IA/SP/2018. Appsilon funded the machine learning and software development, as well as the costs of cloud computing.

## AUTHORS’ CONTRIBUTIONS STATEMENT

KF and RJB conceptualised the research idea, collected parts of the data and prepared annotations. AC, JŚ, JK and PPS were involved in the early stages of the machine learning development, while AC and JŚ completed the training and analysed the results. CP and WCO provided parts of the data. KF, RJB and JŚ led the writing of the manuscript. All authors contributed critically to the drafts and gave final approval for publication.

