## Supplement for "Using machine learning to count Antarctic shag (*Leucocarbo bransfieldensis*) nests on images captured by Remotely Piloted Aircraft Systems"

**SUPLEMENT**

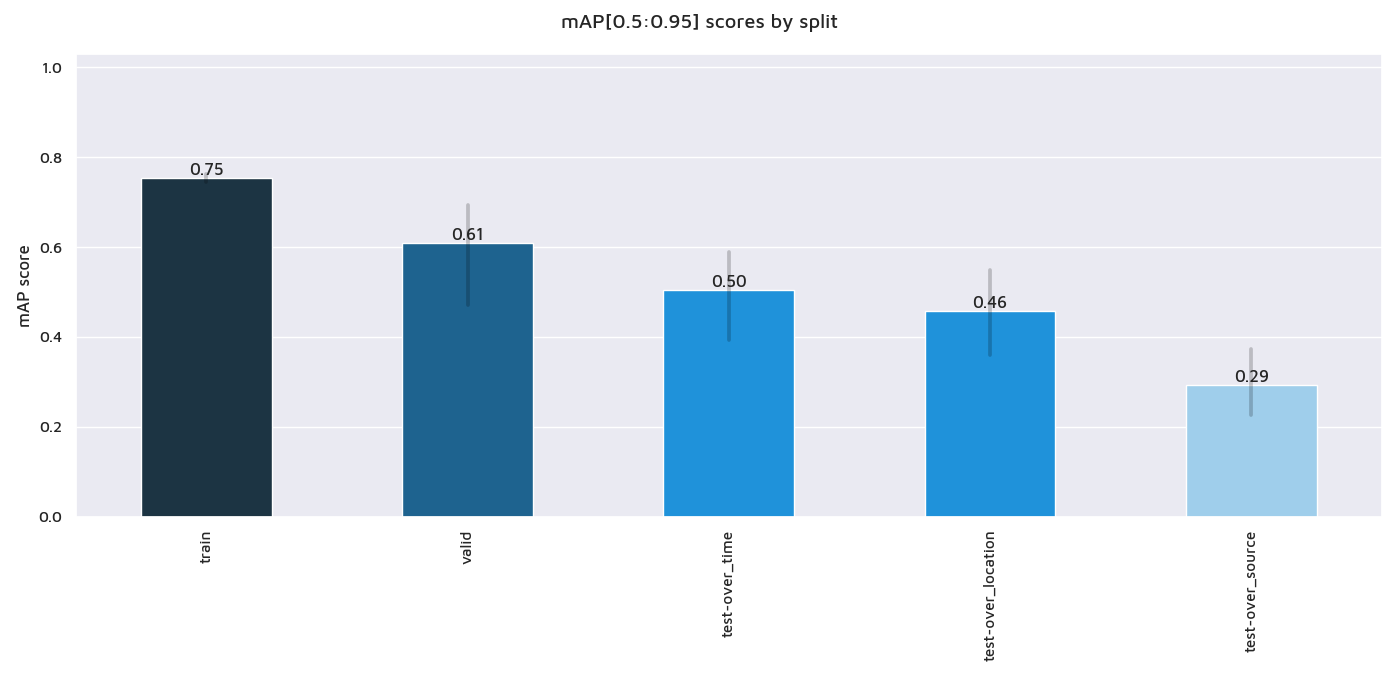

**Fig. S1**. A summary of the mean average precision mAP[0.5:0.95] scores obtained on different parts of the data.

The mean average precision is a standard metric (often referred to as the mAP or mAP[0.5:0.95]) used to assess the performance of object detection models (Eikelboom, et al. 2019, Moreni et al. 2023). mAP[0.5:0.95] means that the average precision AP is calculated for ten intersection over union (IoU) thresholds, from 0.5 to 0.95 with a step of 0.05, and then averaged over the number of classes (Moreni et al. 2023). The IoU is a number from 0 to 1 that specifies the amount of overlap between the predicted and ground truth bounding boxes. In the present study, the mAP score was not an optimal metric. This is mainly due to the spatial precision of the available annotations; while they correctly indicate the presence of a nest in most cases, they often do not exactly match the shape of the nest (this is especially true for the data captured in 2016 and is reflected in the significantly lower mAP scores). For completeness, we present a summary graph with mAP scores (Figure S1) and detailed graphs with each orthomosaic’s score presented separately (Figure S3).

**Table S1.** Detailed view into the data used in the study

| Location and date of capture | resolution [mm/px] | width [m] | length [m] | location (lon) | location (lat) |
| --- | --- | --- | --- | --- | --- |
| Shag Rock 2019-11-26 | 6 | 62 | 43 | -62.18448 | -58.29773 |
| Shag Rock 2019-12-20 | 6 | 68 | 47 | -62.18446 | -58.29781 |
| Shag Rock 2020-01-30 | 6 | 69 | 50 | -62.18444 | -58.29784 |
| Shag Rock 2020-02-22 | 6 | 60 | 43 | -62.18447 | -58.29769 |
| Shag Rock 2020-09-20 | 9 | 134 | 119 | -62.18413 | -58.29853 |
| Shag Rock 2020-10-08 | 6 | 95 | 64 | -62.18433 | -58.29824 |
| Shag Rock 2020-11-15 | 6 | 67 | 46 | -62.18445 | -58.29781 |
| Shag Rock 2020-12-19 | 7 | 66 | 49 | -62.18445 | -58.29777 |
| Shag Rock 2020-12-26 | 7 | 73 | 56 | -62.18441 | -58.29789 |
| Shag Rock 2021-01-12 | 6 | 114 | 88 | -62.1842 | -58.29803 |
| Shag Rock 2021-01-28 | 6 | 72 | 58 | -62.18439 | -58.29789 |
| Shag Rock 2021-02-04 | 6 | 67 | 45 | -62.18447 | -58.29781 |
| Shag Rock 2021-03-26 | 6 | 74 | 59 | -62.1844 | -58.29792 |
| Shag Rock 2021-04-13 | 6 | 70 | 45 | -62.18445 | -58.29785 |
| Shag Rock 2020-01-10 | 7 | 118 | 90 | -62.18415 | -58.29802 |
| Shag Rock 2020-03-17 | 6 | 63 | 44 | -62.18446 | -58.29773 |
| Shag Rock 2020-11-26 | 6 | 70 | 52 | -62.18442 | -58.29784 |
| Shag Rock 2021-02-21 | 7 | 71 | 49 | -62.18444 | -58.29786 |
| Shag Rock 2021-03-10 | 6 | 75 | 54 | -62.18442 | -58.29794 |
| Shag Rock 2021-09-30 | 6 | 65 | 45 | -62.18445 | -58.29777 |
| Shag Rock 2021-11-03 | 7 | 65 | 44 | -62.18447 | -58.29776 |
| Shag Rock 2021-11-24 | 7 | 84 | 63 | -62.18438 | -58.29799 |
| Shag Rock 2021-12-06 | 6 | 50 | 39 | -62.18449 | -58.2975 |
| Shag Rock 2021-12-30 | 6 | 54 | 38 | -62.18449 | -58.29758 |
| Shag Rock 2022-01-05 | 7 | 78 | 76 | -62.18425 | -58.29799 |
| Shag Rock 2022-01-12 | 6 | 68 | 47 | -62.18445 | -58.29782 |
| Shag Rock 2022-01-23 | 6 | 73 | 52 | -62.18444 | -58.29791 |
| Shag Rock 2022-02-09 | 7 | 67 | 46 | -62.18445 | -58.29781 |
| Shag Rock 2022-02-18 | 6 | 64 | 45 | -62.18445 | -58.29779 |
| Shag Rock 2022-02-28 | 6 | 68 | 45 | -62.18446 | -58.29782 |
| Shag Rock 2022-02-28 | 9 | 68 | 46 | -62.18445 | -58.29782 |
| Cape Melville 2022-12-04 | 20 | 336 | 207 | -62.02162 | -57.5832 |
| Cape Melville 2022-12-12 | 18 | 401 | 257 | -62.02157 | -57.58436 |
| Turret Point 2020-12-31 | 14 | 55 | 59 | -62.08852 | -57.95059 |
| Turret Point 2021-11-23 | 11 | 55 | 59 | -62.08852 | -57.95059 |
| Turret Point 2022-01-06 | 11 | 55 | 59 | -62.08852 | -57.95059 |
| Shag Rock 2022-09-24 | 6 | 67 | 47 | -62.18444 | -58.29779 |
| Shag Rock 2022-11-15 | 6 | 67 | 47 | -62.18445 | -58.29779 |
| Turret Point 2022-10-29 | 10 | 57 | 62 | -62.08849 | -57.9506 |
| Turret Point 2022-12-12 | 11 | 76 | 76 | -62.08842 | -57.95083 |
| Cape Melville 2023-11-11 | 20 | 397 | 261 | -62.02162 | -57.5832 |
| Unnamed Island B 2022-12-26 | 12 | 101 | 101 | -62.2451 | -58.49369 |
| Unnamed Island A 2022-12-12 | 12 | 42 | 31 | -62.00157 | -57.60608 |
| Trowbridge Island 2022-12-12 | 12 | 29 | 47 | -61.99394 | -57.63565 |
| Fregata Island 2016-12-11 | 20 | 279 | 178 | -62.22902 | -59.10225 |
| Rzepecki Island 2016-12-28 | 20 | 139 | 104 | -62.09285 | -58.84302 |
| Nelson Island 2018-12-27 | 25 | 143 | 102 | -62.2896 | -59.2315 |
| Rzepecki Island 2016-12-30 | 20 | 139 | 104 | -62.09285 | -58.84302 |
| Kwarecki Island 2016-12-28 | 34 | 182 | 136 | -62.11832 | -58.8915 |
| Unnamed Island C 2016-12-25 | 15 | 157 | 113 | -62.19369 | -59.05515 |

**Table S2.** Detailed performance of the model

| Location and date of capture | Split | F1 score | mAP[0.5:0.95] | n_nests | n_preds | TP | FP | FN |
| --- | --- | --- | --- | --- | --- | --- | --- | --- |
| Cape Melville 2022-12-04 | train | 0.997 | 0.786 | 459 | 460 | 458 | 2 | 1 |
| Cape Melville 2022-12-12 | train | 0.998 | 0.766 | 459 | 461 | 459 | 2 | 0 |
| Shag Rock 2019-11-26 | train | 1 | 0.8 | 69 | 69 | 69 | 0 | 0 |
| Shag Rock 2019-12-20 | train | 1 | 0.746 | 62 | 62 | 62 | 0 | 0 |
| Shag Rock 2020-01-30 | train | 0.992 | 0.703 | 59 | 60 | 59 | 1 | 0 |
| Shag Rock 2020-02-22 | train | 1 | 0.694 | 60 | 60 | 60 | 0 | 0 |
| Shag Rock 2020-09-20 | train | 0.991 | 0.762 | 54 | 55 | 54 | 1 | 0 |
| Shag Rock 2020-10-08 | train | 1 | 0.745 | 57 | 57 | 57 | 0 | 0 |
| Shag Rock 2020-11-15 | train | 1 | 0.788 | 74 | 74 | 74 | 0 | 0 |
| Shag Rock 2020-12-19 | train | 1 | 0.779 | 71 | 71 | 71 | 0 | 0 |
| Shag Rock 2020-12-26 | train | 1 | 0.764 | 71 | 71 | 71 | 0 | 0 |
| Shag Rock 2021-01-12 | train | 0.993 | 0.745 | 70 | 71 | 70 | 1 | 0 |
| Shag Rock 2021-01-28 | train | 1 | 0.735 | 66 | 66 | 66 | 0 | 0 |
| Shag Rock 2021-02-04 | train | 0.992 | 0.746 | 62 | 63 | 62 | 1 | 0 |
| Shag Rock 2021-03-26 | train | 1 | 0.791 | 65 | 65 | 65 | 0 | 0 |
| Shag Rock 2021-04-13 | train | 1 | 0.768 | 65 | 65 | 65 | 0 | 0 |
| Shag Rock 2020-01-10 | train | 1 | 0.763 | 58 | 58 | 58 | 0 | 0 |
| Shag Rock 2020-03-17 | train | 1 | 0.7 | 59 | 59 | 59 | 0 | 0 |
| Shag Rock 2020-11-26 | train | 1 | 0.807 | 75 | 75 | 75 | 0 | 0 |
| Shag Rock 2021-02-21 | train | 1 | 0.739 | 68 | 68 | 68 | 0 | 0 |
| Shag Rock 2021-03-10 | train | 1 | 0.764 | 72 | 72 | 72 | 0 | 0 |
| Shag Rock 2021-09-30 | train | 1 | 0.772 | 79 | 79 | 79 | 0 | 0 |
| Shag Rock 2021-11-03 | train | 0.994 | 0.768 | 81 | 82 | 81 | 1 | 0 |
| Shag Rock 2021-11-24 | train | 1 | 0.771 | 80 | 80 | 80 | 0 | 0 |
| Shag Rock 2021-12-06 | train | 1 | 0.776 | 80 | 80 | 80 | 0 | 0 |
| Shag Rock 2021-12-30 | train | 1 | 0.762 | 77 | 77 | 77 | 0 | 0 |
| Shag Rock 2022-01-05 | train | 1 | 0.758 | 75 | 75 | 75 | 0 | 0 |
| Shag Rock 2022-01-12 | train | 1 | 0.729 | 76 | 76 | 76 | 0 | 0 |
| Shag Rock 2022-01-23 | train | 1 | 0.731 | 77 | 77 | 77 | 0 | 0 |
| Shag Rock 2022-02-09 | train | 1 | 0.722 | 69 | 69 | 69 | 0 | 0 |
| Shag Rock 2022-02-18 | train | 1 | 0.722 | 74 | 74 | 74 | 0 | 0 |
| Shag Rock 2022-02-28 | train | 0.993 | 0.714 | 69 | 68 | 68 | 0 | 1 |
| Shag Rock 2022-02-28 | train | 1 | 0.77 | 69 | 69 | 69 | 0 | 0 |
| Turret Point 2020-12-31 | valid | 0.955 | 0.635 | 70 | 64 | 64 | 0 | 6 |
| Turret Point 2021-11-23 | valid | 1 | 0.723 | 67 | 67 | 67 | 0 | 0 |
| Turret Point 2022-01-06 | valid | 0.934 | 0.472 | 63 | 59 | 57 | 2 | 6 |
| Shag Rock 2022-09-24 | test-over_time | 0.993 | 0.553 | 73 | 74 | 73 | 1 | 0 |
| Shag Rock 2022-11-15 | test-over_time | 0.981 | 0.513 | 79 | 82 | 79 | 3 | 0 |
| Turret Point 2022-10-29 | test-over_time | 0.991 | 0.642 | 58 | 59 | 58 | 1 | 0 |
| Turret Point 2022-12-12 | test-over_time | 0.974 | 0.505 | 59 | 56 | 56 | 0 | 3 |
| Cape Melville 2023-11-11 | test-over_time | 0.946 | 0.312 | 478 | 482 | 454 | 28 | 24 |
| Unnamed Island B 2022-12-26 | test-over_location | 1 | 0.36 | 9 | 9 | 9 | 0 | 0 |
| Unnamed Island A 2022-12-12 | test-over_location | 0.985 | 0.549 | 32 | 33 | 32 | 1 | 0 |
| Trowbridge Island 2022-12-12 | test-over_location | 1 | 0.467 | 7 | 7 | 7 | 0 | 0 |
| Nelson Island 2018-12-27 | test-over_source | 0.949 | 0.485 | 72 | 65 | 65 | 0 | 7 |
| Fregata Island 2016-12-20 | test-over_source | 0.65 | 0.189 | 27 | 13 | 13 | 0 | 14 |
| Kwarecki Island 2016-12-28 | test-over_source | 0.909 | 0.211 | 6 | 5 | 5 | 0 | 1 |
| Rzepecki Island 2016-12-30 | test-over_source | 0.967 | 0.332 | 62 | 58 | 58 | 0 | 4 |
| Unnamed Island C 2016-12-25 | test-over_source | 0.842 | 0.317 | 11 | 8 | 8 | 0 | 3 |
| Fregata Island 2016-12-11 | test-over_source | 0.926 | 0.328 | 29 | 25 | 25 | 0 | 4 |
| Rzepecki Island 2016-12-28 | test-over_source | 0.615 | 0.184 | 63 | 28 | 28 | 0 | 35 |

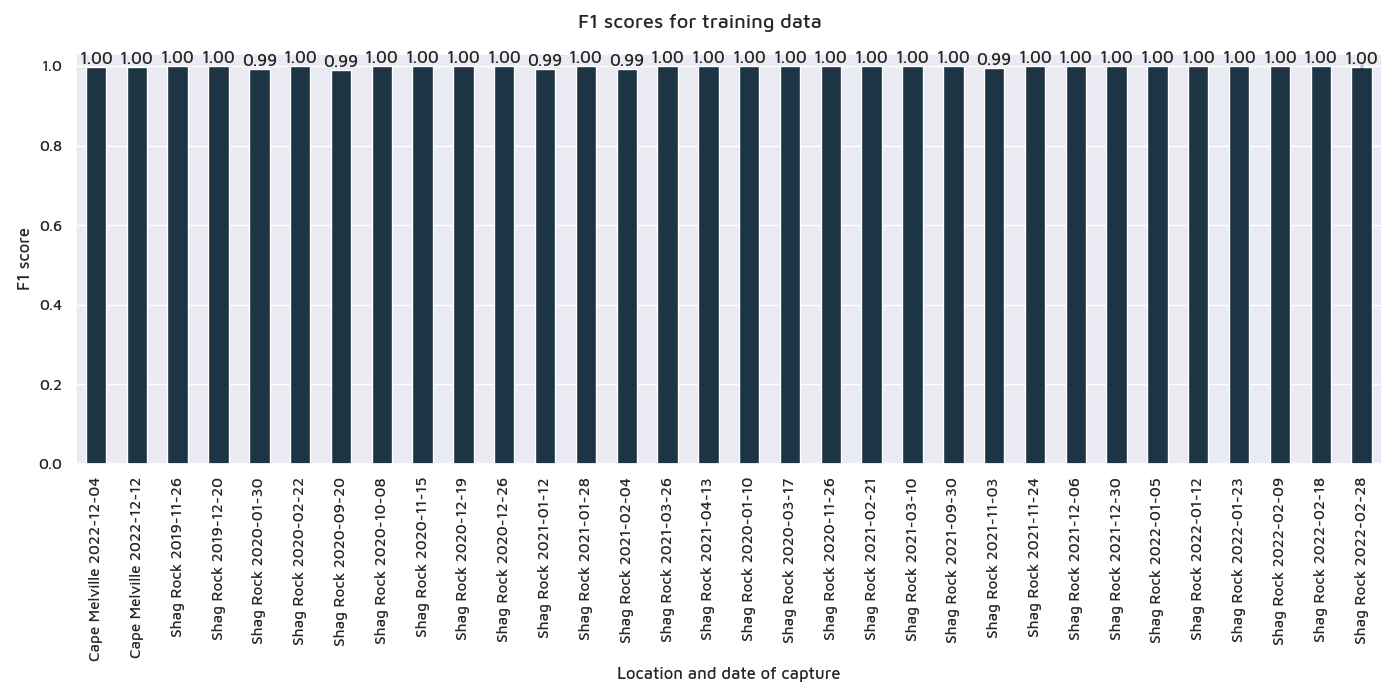

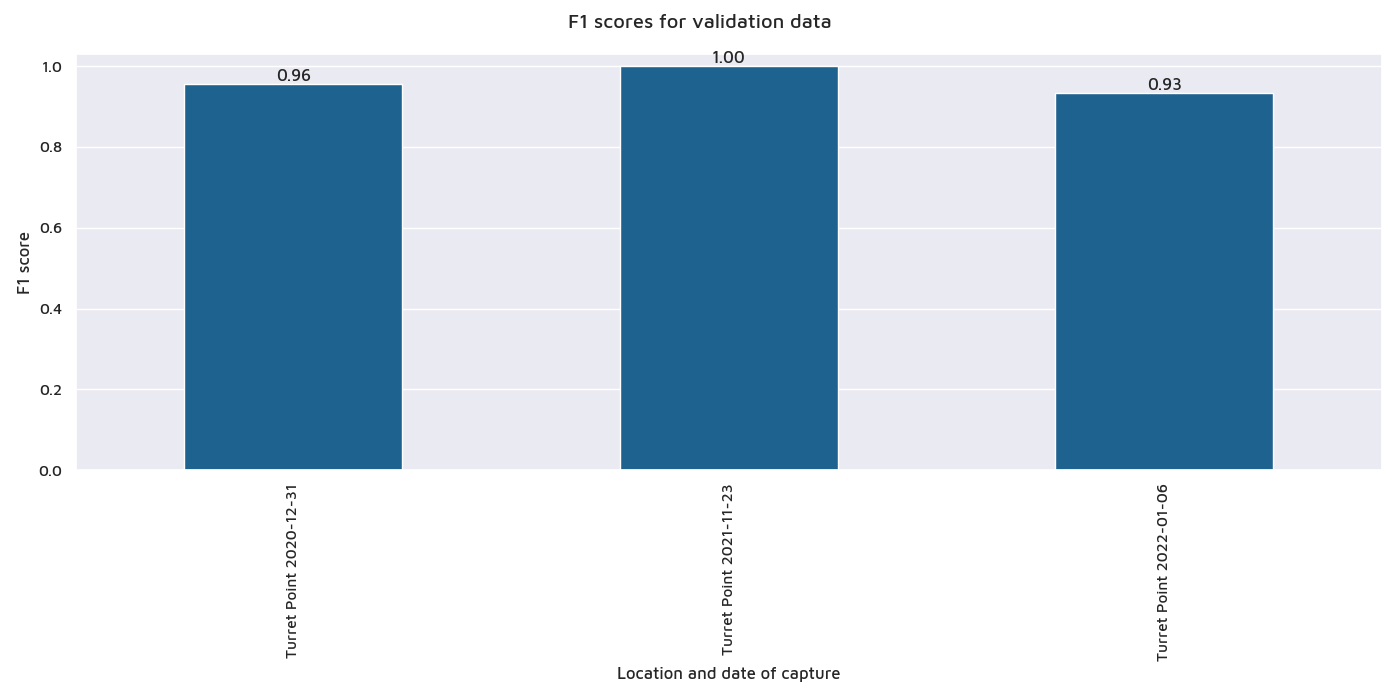

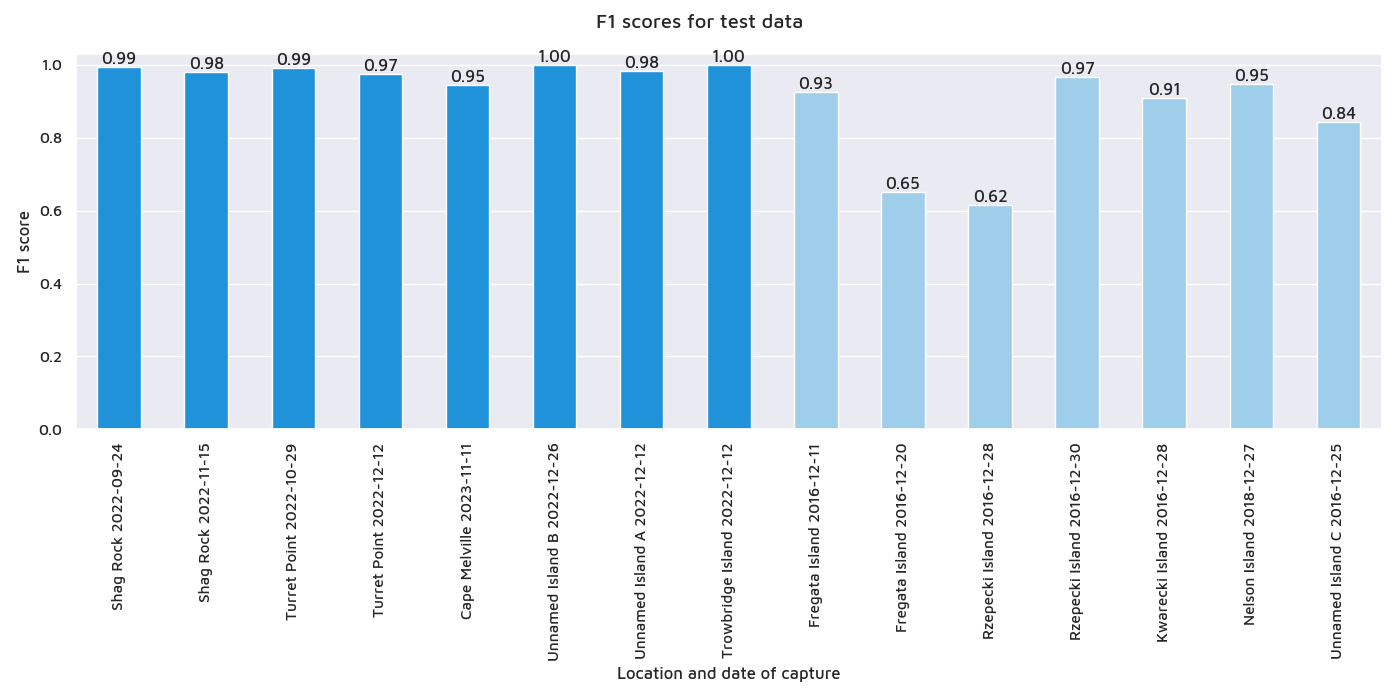

**Fig. S2.** Detailed F1 scores of the model on training, validation and test data

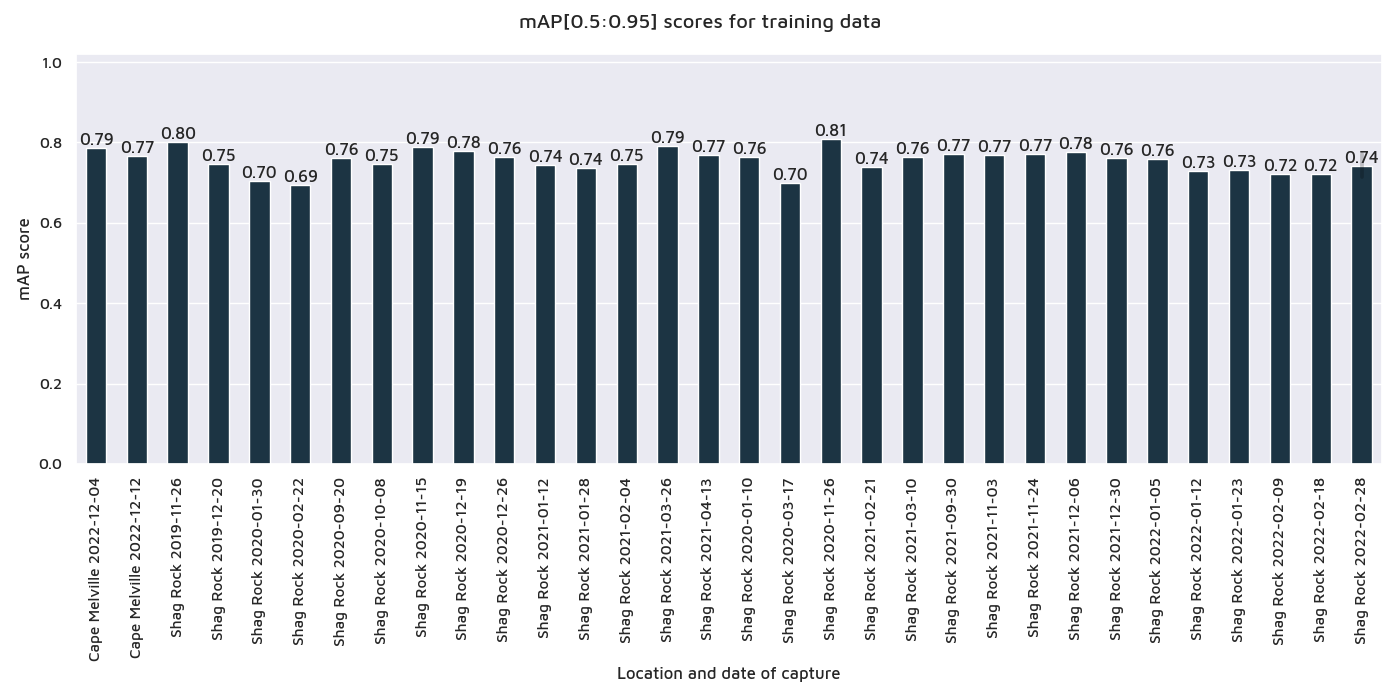

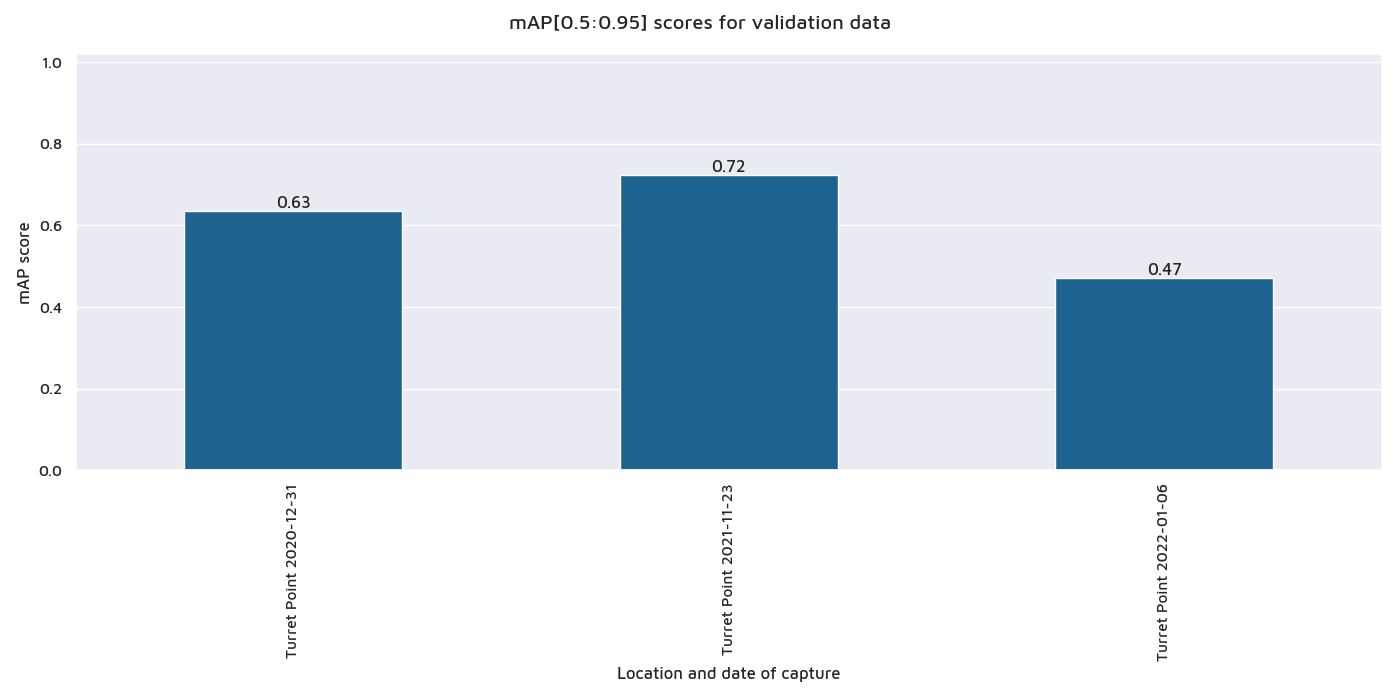

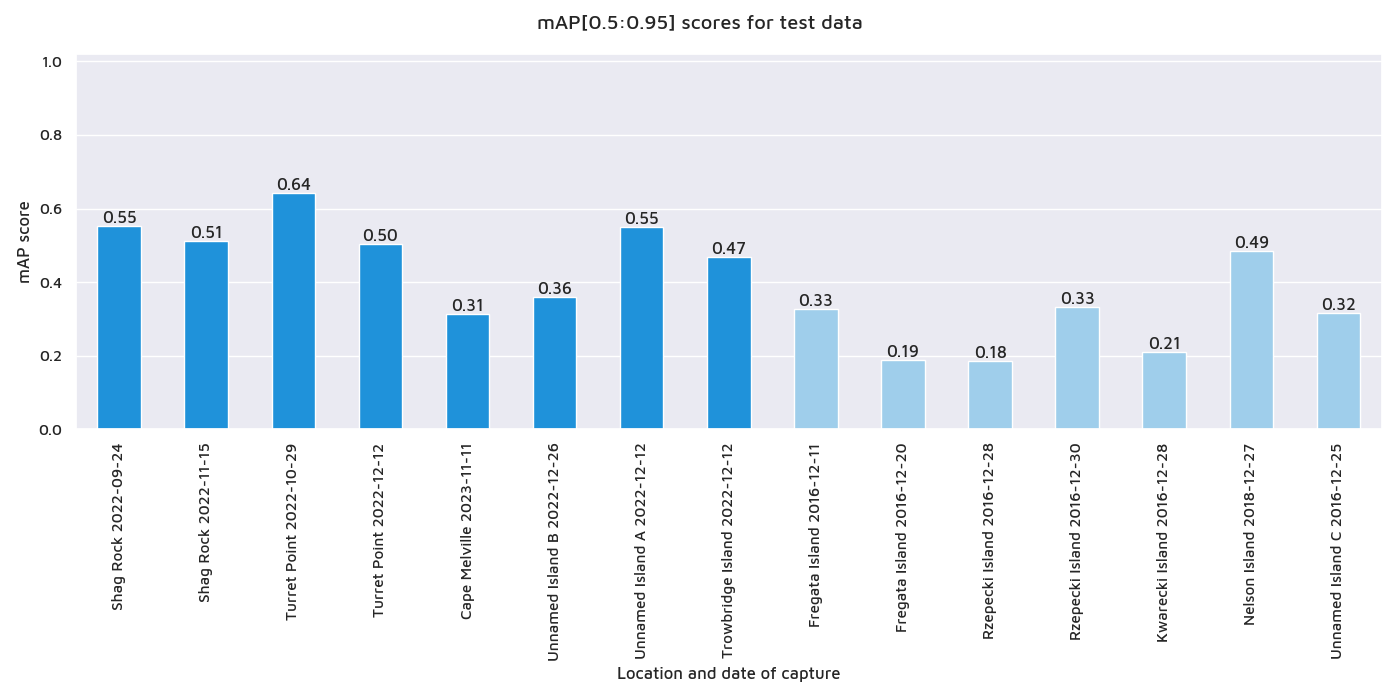

**Fig. S3**. Detailed mAP[0.5:0.95] scores for the individual images
